# Discovering Governing Equations of Biological Systems through Representation Learning and Sparse Model Discovery

**DOI:** 10.1101/2024.09.19.613953

**Authors:** Mehrshad Sadria, Vasu Swaroop

**Affiliations:** Department of Applied Mathematics, University of Waterloo, Waterloo, Ontario N2L 3G1, Canada; Department of Computer Science Information Systems, BITS-Pilani, Pilani Campus, Pilani, 333031, India

## Abstract

Understanding the governing rules of complex biological systems remains a significant challenge due to the nonlinear, high-dimensional nature of biological data. In this study, we present CLERA, a novel end-to-end computational framework designed to uncover parsimonious dynamical models and identify active gene programs from single-cell RNA sequencing data. By integrating a supervised autoencoder architecture with Sparse Identification of Nonlinear Dynamics, CLERA leverages prior knowledge to simultaneously extract related low-dimensional embeddings and uncovers the underlying dynamical systems that drive the processes. Through the analysis of both synthetic and biological datasets, CLERA demonstrates robust performance in reconstructing gene expression dynamics, identifying key regulatory genes, and capturing temporal patterns across distinct cell types. CLERA’s ability to generate dynamic interaction networks, combined with network rewiring using Personalized PageRank to highlight central genes and active gene programs, offers new insights into the complex regulatory mechanisms underlying cellular processes.

## Introduction

Across many scientific disciplines, discovering governing equations has traditionally served as the cornerstone of understanding systems (1). Derived from mathematical and physical laws, these equations provide interpretable and generalizable frameworks for explaining and predicting various phenomena. In areas such as biology (2), epidemiology (3), and finance (4), mathematical models are used to model signalling pathways, population dynamics and disease spread, and market fluctuations, respectively. However, for complex systems with high dimensionality and nonlinearity, including biological processes, traditional approaches often fall short (5). Discovering the main equations governing these systems can be challenging, and even when partial knowledge exists, relying solely on first principles becomes impractical (6).

The modern era, with its abundance of data and computational power, has facilitated the emergence of data-driven model discovery as a powerful paradigm in scientific exploration (7). This approach directly leverages data to uncover the hidden principles that govern complex systems. In the context of cellular biology, single-cell RNA sequencing (scRNA-seq) provides an unprecedented window into individual cells, which offers insights into gene expression variation across diverse cellular populations (8). By analyzing this data, researchers can investigate the molecular machinery underlying development, disease, and response to external perturbation (9,10). The noisy, nonlinear, and high-dimensional characteristics of scRNA-seq data and the biological processes it captures pose significant challenges for analysis and interpretation (11). These complexities make it difficult to uncover the underlying principles of biological processes and pinpoint their key drivers (12). While previous methods have achieved success in specific tasks, limitations remain. For instance, the correlative nature of most methods prevents them from capturing causal features and true representations, thus limiting their generalizability (13). Furthermore, these models struggle to discover governing relationships among underlying variables in a parsimonious manner, similar to classical physics settings, which further hinders true interpretability. Therefore, a crucial step in understanding any biological process lies in developing models that not only can accurately predict but also reveal the underlying connections between features in an interpretable and parsimonious manner (14). This ensures the models can be applied across diverse environments and provides clearer insights into the mechanisms governing the process (1).

In the realm of high-dimensional biological data like scRNA-seq, the ability to capture causal representations of the data becomes particularly valuable. This approach goes beyond identifying correlations and allows us to understand the true relationships between variables (15). In this context, identifiability, the ability to uniquely recover the underlying causal structure from observed data, becomes a crucial aspect of representation learning. Traditional methods like Independent Component Analysis (ICA) have achieved success in many areas of linear representation learning (16). In fact, if all latent components are non-Gaussian and independent, ICA can be identifiable. However, ICA struggles with the inherent nonlinearities and complex interactions present in biological data (16). While perfect identifiability, especially in non-linear settings, remains a challenge, incorporating temporal structure, employing additional tasks, or using auxiliary information can facilitate the way to attain identifiability (17,18). Notably, autoencoders can offer a promising avenue for achieving identifiability in non-linear settings (19). By carefully designing their architecture and loss function, autoencoders can help extract meaningful representations from complex biological data (20–23).

In this work, we present CLERA (Cellular Latent Equation and Representation Analysis), a novel end-to-end computational framework that combines the power of data-driven model discovery, specifically Sparse Identification of Nonlinear Dynamics (SINDy), and representation learning. Leveraging a supervised autoencoder architecture, CLERA simultaneously extracts a compact and relevant representation from high-dimensional data and uses it to discover the underlying low-dimensional, non-linear dynamical model governing the system. This learned embedding further allows us to not only identify active gene programs and key genes but also track their transitions over time across cell types, providing insights into the complex dynamic regulatory mechanisms of biological systems. We validate CLERA’s performance on both simulated data (with known active gene programs) with different sizes and real-world biological datasets.

## Result

### Discovery of Dynamical Systems and Gene Programs from Simulated Data

We first investigate the performance of the SINDy part of CLERA in discovering the underlying governing equations of a simple simulated biological system with two driver genes (Fig. 1a). The dynamics of this system are described by a well-established set of differential equations commonly used in various biological contexts such as the lac operon, metabolic signalling pathways, and the cell cycle (24). Synthetic data is generated using this system of equations with varying noise levels. We then apply SINDy, to recover the equations. Notably, the governing equations are discovered with high accuracy. Figure 1a shows this successful reconstruction, with the recovered parameters closely mirroring the original values (Method section). Also, the results from solving the discovered differential equation closely match the generated data, further validating the accuracy of the equations discovered (Fig. 1b).

**Figure 1.**
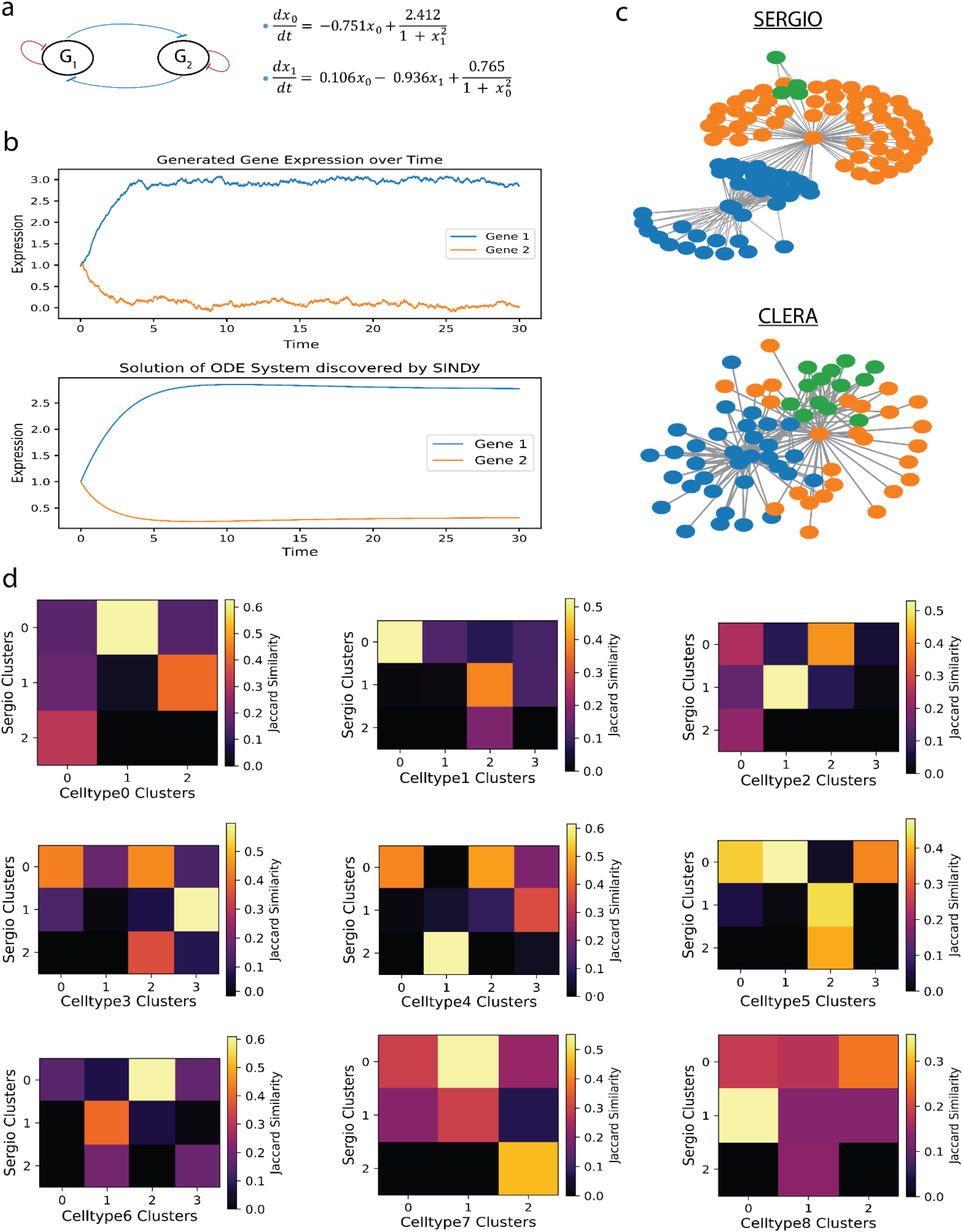
CLERA discovers dynamical systems and gene programs from simulated data. a, Schematic of a two-gene regulatory network (G_1_ and G_2_) with discovered governing equations and parameters shown. b, Comparison of generated gene expression data (top) and solutions from equations discovered by SINDy (bottom) for the two-gene system over time. Gene 1 and Gene 2 expression levels are plotted against time. c, Gene interaction networks for cell type 7 derived from SERGIO ground truth (top) and CLERA (bottom). Nodes represent genes, coloured by gene programs identified through clustering. d, Heatmaps showing Jaccard similarity between SERGIO and CLERA-derived gene program clusters across nine cell types (CellType0 to CellType8). Colour intensity indicates the degree of similarity, with lighter colours representing higher similarity and darker colours lower similarity.

Then to evaluate the performance of CLERA in a more realistic scenario with a larger dataset, we apply it to simulated data generated by SERGIO, which incorporates various types of noise for realistic data generation (25). We train CLERA on a simulated dataset, with 6300 cells across 100 genes and nine distinct cell types. We hypothesize that an optimal representation learned by the model should not only achieve high accuracy in data reconstruction (autoencoder loss) but also should perform well in tasks such as cell type classification and sparse dynamical model discovery.

To address stochasticity and ensure robustness in finding the optimal latent embedding, we run CLERA multiple times (50 for this data) using various initial conditions. We select the model with the lowest combined loss (method section) while also prioritizing parsimony in the discovered model. In our analysis, we observe that CLERA successfully identified a latent embedding with high accuracy in both reconstruction and classification (Supplementary Figure 1, Supplementary Figure 5a).

We then leverage the representation learned by CLERA to identify active gene programs and their dynamics over time. To uncover the connection between latent nodes and genes we compute SHAP (26) values between each node in the autoencoder’s latent layer and genes and rank the identified genes based on their SHAP values (choosing the top 30 genes for each node). Using the results from the SHAP method and the discovered differential equations, we construct a network of interactions between latent nodes and genes. We then apply Personalized PageRank (PPR) to this network, starting from each gene, to identify the most relevant genes for the selected gene (27). This approach enables us to refine the network by selecting only the top connected genes with the highest PPR scores while filtering out the latent nodes. A clustering algorithm is applied to this graph to detect the gene programs. Given that CLERA can uniquely incorporate a time component, this process can be done for different stages of the trajectory and cell types. To assess CLERA’s performance in capturing active gene programs, we perform the same clustering analysis on the SERGIO ground truth network and compare the resulting clusters obtained (Fig. 1c for celltype7). We also observed a high degree of similarity between the gene programs of the SERGIO predefined network and the identified gene interaction networks for each cell type, as measured by the Jaccard similarity. This suggests that the latent embedding learned by CLERA can effectively capture active gene programs (Fig. 1d).

### CLERA Uncovers Dynamics and Gene Programs in Pancreatic Development

We further evaluate CLERA on biological scRNA-seq data from mouse pancreas during embryonic development. This dataset comprises 3696 cells, with 27998 genes clustered into eight distinct cell types (28). Following hyperparameter optimization and preprocessing, we trained CLERA several times with varying initializations (Methods), using the gene expression and computed pseudotime (cell ordering) information. CLERA successfully identifies a set of sparse and interpretable differential equations with all individual loss terms in our total loss function decreasing (Fig 2a, Supplementary Figure 2). Also, we observe the temporal dynamic of different latent variables captures distinct patterns for each cell type (Fig. 2b, Supplementary Figure 9a). CLERA also achieves a high classification accuracy using the latent variables where certain latent variables emerge as dominant predictors for individual cell types (Supplementary Figure 5b, Supplementary Figure 7).

**Figure 2.**
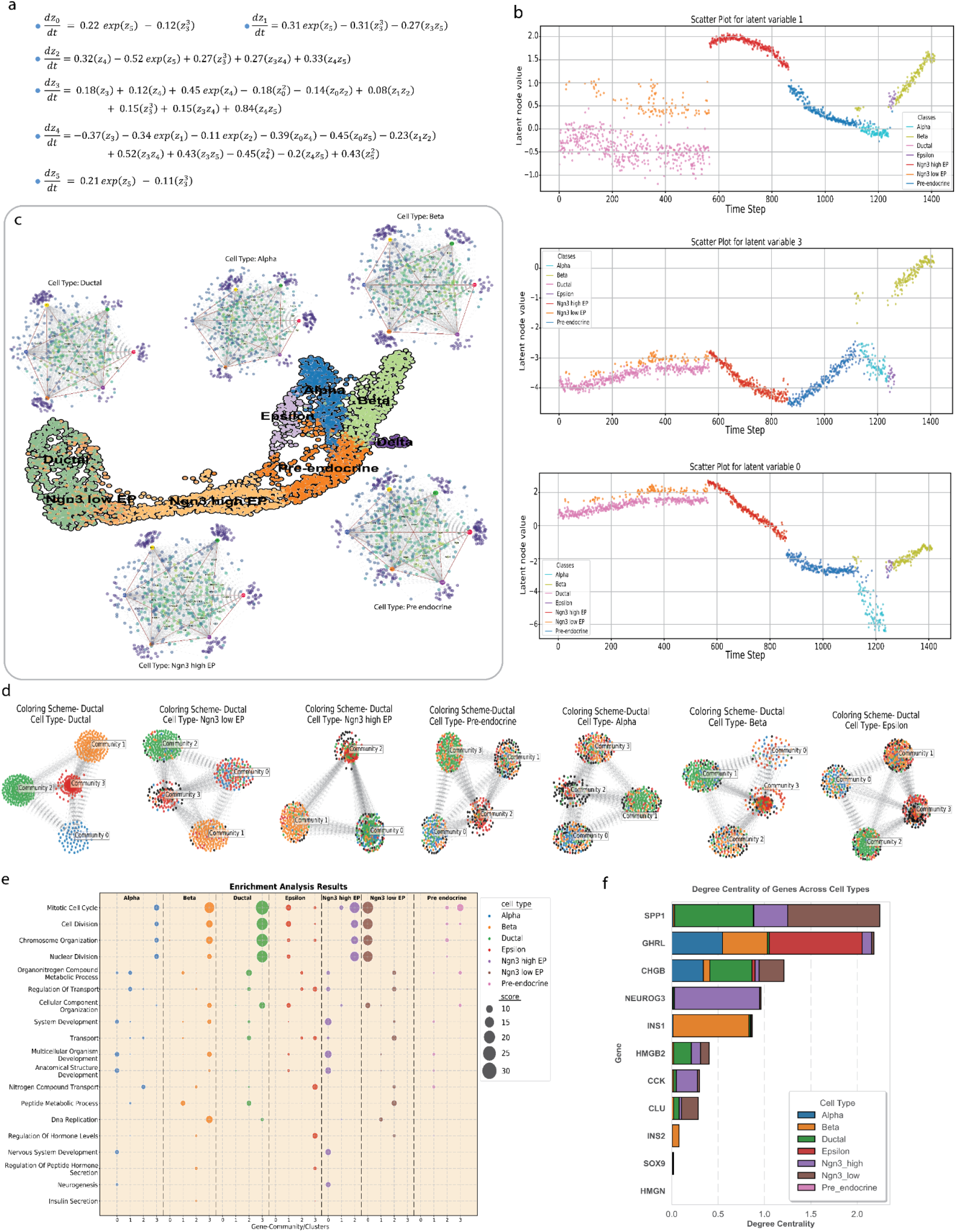
CLERA uncovers dynamics and gene programs in pancreatic development. a, Discovered differential equations governing mouse pancreas development data from scRNA-seq, showing sparse and interpretable models and connections between latent variables. b, Temporal dynamics of latent variables, which illustrate distinct patterns across cell types. c, Interaction graphs for various stages of pancreatic development, show dynamic gene interactions over time for different cell types. d, Clustering results showing gene program similarities across cell types, with shared genes in cluster 1 among Ductal, Ngn3 low EP, and Ngn3 high EP cell types. e, Gene Set Enrichment Analysis results indicating pathways related to pancreatic development and neurogenesis. f, Degree centrality analysis identifying key genes for each cell type, including Spp1, Chgb, Neurog3, Ins1, Ins2, Clu, and Sox9.

To explore the connection between latent nodes and genes, we calculate the SHAP values (29) for each gene-latent node pair and identify the top “K” genes (K=300 for this data) connected to each latent node, ranked by their absolute values. Using the discovered equations (latent node-latent node interaction) and SHAP values (gene-latent node interaction), we generate a series of interaction graphs for various stages of pancreas development (Fig. 2c). Unlike traditional network inference methods, which only produce a static graph for the whole process, our approach captures dynamic graphs over time. Next, we apply a clustering algorithm to the interaction graphs to identify groups of interconnected and potentially co-regulated genes. These graphs, representing different stages of pancreatic development, allowed us to observe changes in gene interactions over time. To understand the similarity of active gene programs across different cell types, we analyze the clustering results for a specific cell type and transfer the identified gene colours to the analysis of other cell types (Fig. 2d). We observe a high degree of similarity in shared genes for cluster 1 among Ductal, Ngn3 low EP, and Ngn3 high EP cell types. Analyzing these shared genes reveals several previously known key genes, such as Sox9, Neurog3, Hes1, Foxa3, and Nfib, as well as important signalling pathways like Wnt, Notch, and TGF-β (30). Furthermore, Gene Set Enrichment Analysis reveals several pathways related to pancreatic development, demonstrating the biological relevance of the gene programs discovered by CLERA (Fig. 2e). Interestingly, CLERA also captures pathways involved in neurogenesis and neural development, which aligns with previous studies and highlights the molecular and cellular similarities between pancreatic and neural cell differentiation (28,31).

To identify key and central genes for each cell type, we restructure the interaction network using the PPR technique to remove latent nodes. This network rewiring allows us to focus directly on gene interactions. We then apply centrality measures to the restructured network to identify the most influential genes for each cell type (Fig. 2f, Supplementary Figure 10a). As a result of the centrality analysis, several key genes are identified, including Spp1 (32), a regulator of the epithelial-mesenchymal transitory axis and duct cell de-differentiation; Chgb (33), a neuroendocrine cell marker; and Neurog3 (34,35), crucial for endocrine cell differentiation. The analysis also confirms the central roles of Ins1 and Ins2 in beta cell function, along with Clu (36) and Sox9 (37), both critical for progenitor cell maintenance and differentiation. CLERA correctly captures these key genes, aligning with prior studies that emphasize their importance in pancreatic development.

### CLERA Reveals Central Genes and Dynamics in Hematopoietic Differentiation

Next, we investigate bone marrow development, examining the differentiation of hematopoietic stem cells (HSPCs) into erythroids, monocytes, and dendritic cells (DCs). This dataset comprises 5780 cells and 14319 genes clustered into 10 distinct cell types (38).

To enhance CLERA’s performance on this data, we apply transfer learning from our previous pancreas study (method section). By initializing CLERA with pre-trained weights, we leverage the knowledge and relationships obtained from the previous part, which results in a faster optimization and more accurate representation of the data. Also, we observe a decrease across all components of the loss function, showing that all loss terms were effectively optimized and also parsimonious discovered equations (Fig. 3a, Supplementary Figure 3). Furthermore, investigating the latent space shows that the temporal dynamics of different latent variables capture distinct patterns for each cell type, which shows that the embedding learnt by CLERA can identify and characterize unique behavioural signatures for each cell type (Fig. 3b, Supplementary Figure 9b). We observe that the latent embedding discovered by CLERA achieves high classification accuracy and also shows distinctive cell type-level differentiation, where specific latent variables drive the classification of particular cell types (Supplementary Figure 5c, Supplementary Figure 8).

**Figure 3.**
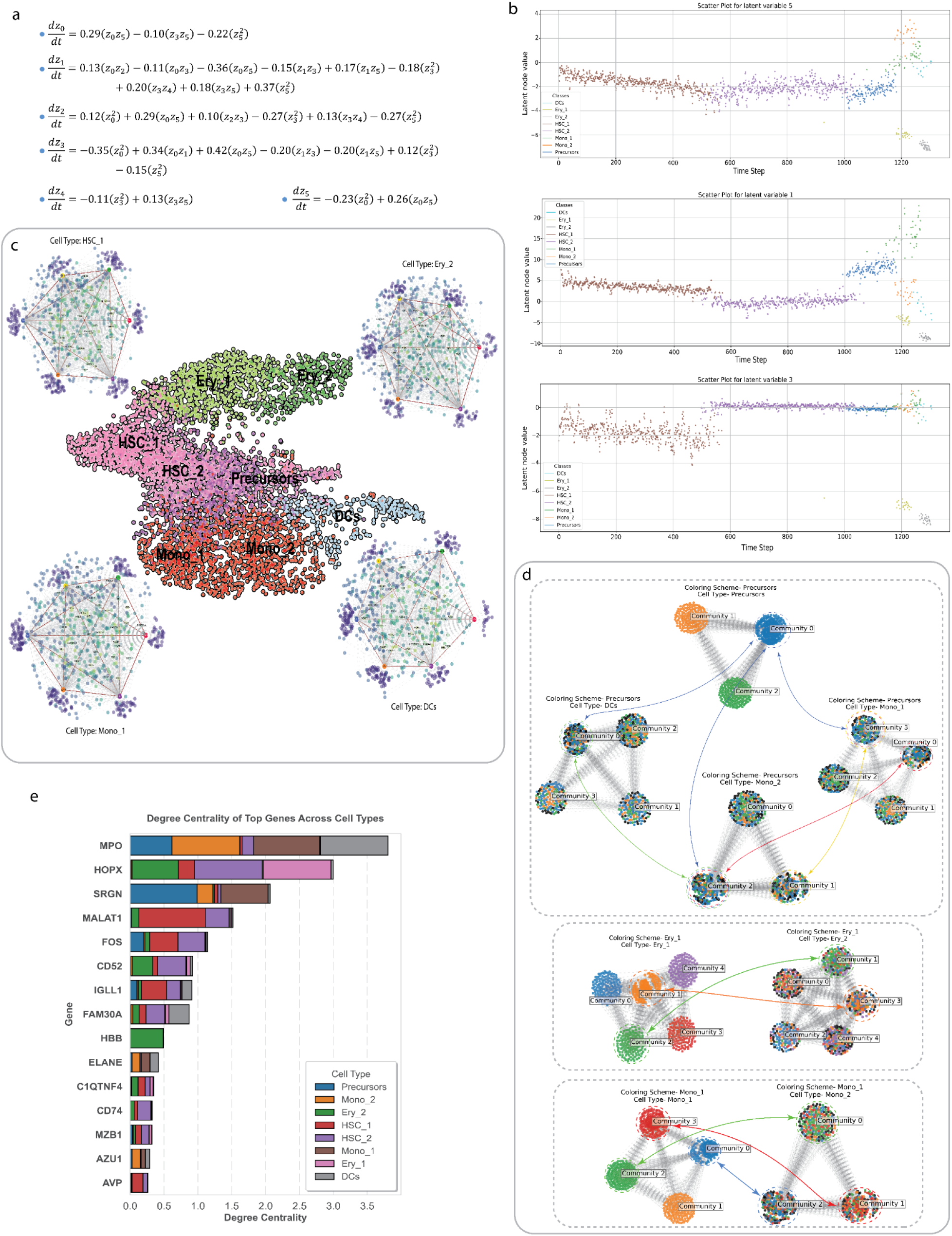
CLERA reveals central genes and dynamics in hematopoietic differentiation. a, Differential equations discovered for bone marrow development data, showing connections between latent variables. b, Temporal dynamics of latent variables, with distinct patterns across cell types. c, Interaction graphs for different stages of bone marrow development, which capture dynamic gene interactions for each cell type. d, Clustering analysis of co-regulated gene groups at each developmental stage, with significant similarities between precursors, monocytes, DCs, and among Ery-1 and Ery-2, Mono-1 and Mono-2 subpopulations. e, Degree centrality analysis identifies key genes driving cellular differentiation, including Mpo, HOPX, Malat1, FOS, CD52, FAM30A, and CD74.

Then, by identifying top genes using the SHAP method and leveraging the discovered equations, we generate a series of graphs representing different stages of bone marrow development (Fig. 3c). Through clustering analysis on these graphs, we identify groups of co-regulated genes at each developmental stage (Fig. 3d). Moreover, using label transfer techniques, we identify a significant similarity in co-regulated genes between precursors (cluster 1), monocytes (cluster 3 in Mono_1 and cluster 2 in Mono_2) and DCs (cluster 0). Some of the key genes discovered have been shown to be crucial for monocyte development, including ID2, TYROBP, FLT3, PDE4B, and GLIPR1. We also observe a strong similarity between the two erythroid subpopulations Ery_1 and Ery_2, particularly between clusters 1 and 3, and between clusters 2 and 1. Similarly, the monocyte subpopulations Mono_1 and Mono_2 show considerable overlap, with cluster 3 in Mono_1 closely aligning with cluster 1 in Mono_2, cluster 2 in Mono_1 resembling cluster 0 in Mono_2, and cluster 0 in Mono_1 closely matching cluster 2 in Mono_2. This suggests that these subpopulations have many common genes, which shows similarities in their developmental pathways and active gene programs.

To identify the critical genes within these networks, we apply PPR for network rewiring, which allows us to remove latent nodes and focus on direct gene interactions. Centrality measures then pinpoint key genes driving cellular differentiation during hematopoiesis (Fig. 3e, Supplementary Figure 10b). MPO (39), crucial for neutrophil differentiation, is identified as a key myeloid marker, while HOPX (39) emerges as a regulator of primitive hematopoiesis, guiding early progenitor cell fate. Malat1 (40), known for regulating gene expression in hematopoietic stem cells, and FOS (41,42), linked to cell proliferation and differentiation under cytokine signalling, are also highlighted. Also, CD52 (43), a marker of mature lymphocytes, FAm30A, which has shown links to immune response regulation and other hematopoietic lineages, and CD74 (44), essential for antigen presentation in immune cells, are captured. CLERA effectively identified these genes, which align with their known roles in hematopoiesis.

## Discussion

Here we introduce a novel method, CLERA which represents a significant advancement in our ability to uncover the underlying principles governing complex biological systems. CLERA integrates data-driven model discovery and representation learning, to provide a robust way for uncovering interpretable and parsimonious dynamical models from high-dimensional scRNA-seq data. The ability to simultaneously learn coordinates and governing equations is perhaps the most complementary approach to traditional physics modelling. While classical methods rely on predefined equations based on known physical laws, CLERA also allows the data to guide the discovery of both the relevant coordinates (latent representation) and the equations that govern their dynamics. This approach is particularly valuable in biological systems where the underlying mechanisms are often complex and not fully understood.

Our results demonstrate CLERA’s ability to accurately recover governing equations from both simulated and real-world biological data. In the simple simulated biological system, CLERA precisely predicts the governing equations, even under varying noise levels. This performance is further validated with a larger simulated dataset generated by SERGIO, which incorporates realistic noise patterns, showing CLERA’s accuracy in modelling complex, high-dimensional data. When applied to real-world datasets from the pancreas and bone marrow development, CLERA leads to biologically relevant insights. The model effectively identifies key genes through network rewiring and centrality measures, as well as detecting active gene programs involved in these processes aligned with previous experimental studies. Another key strength of CLERA lies in its ability to capture the temporal dynamics of gene programs. Unlike traditional network inference methods that produce static graphs, CLERA generates dynamic interaction networks that evolve over time across cell types. This temporal aspect is crucial for understanding differences in complex regulatory mechanisms of cellular processes such as differentiation and development. The identification of potentially novel key genes and gene programs, along with their temporal progression, can present new opportunities for further experimental exploration of dynamic cellular processes.

CLERA integrates multiple components in its loss function to address the challenges of unsupervised learning of identifiable nonlinear representations. The combination of reconstruction loss, classification loss, and SINDy losses introduces inductive biases that guide the model toward learning more meaningful and interpretable embeddings. This comprehensive loss function imposes constraints on latent dynamics, to ensure consistency with known biological principles while maintaining the flexibility to discover new relationships from the data. By balancing these different loss components, CLERA achieves a robust and parsimonious framework for uncovering interpretable dynamical models. Also, the use of transfer learning from the pancreas to the bone marrow dataset demonstrates the model’s adaptability and potential for generalization across different biological contexts. This approach not only improves performance but also suggests that certain learned representations may be conserved across different developmental processes.

In CLERA, we intentionally avoid assuming independence between latent variables, diverging from some recent methods (45). In biological systems, the strict assumption of independence can be problematic, as many genes participate in multiple programs simultaneously. This interconnectedness makes it challenging to capture the true nature of these systems with models that enforce strict independence. By allowing for overlapping gene programs and complex interactions, CLERA’s approach is better suited to reflect the complexity and reality we have in biology. We believe by not imposing independence between latent variables, CLERA can more accurately model the temporal dynamics and complex interactions captured by ODEs, leading to a better understanding of the underlying biological processes.

A critical aspect of using SINDy for Ordinary Differential Equation (ODE) discovery in biological systems is the question of whether there is a single unique way to represent (model) a given process or if multiple valid representations exist. While CLERA effectively recovers governing equations from data, it’s important to recognize that the method may yield different but equally valid solutions due to its consideration of linear transformations and rotations of the underlying dynamical systems. This inherent flexibility reflects the possibility that processes might not have a singular unique representation, but rather several that are mathematically equivalent yet distinct in form. This raises an important challenge in interpreting the biological significance of the discovered ODEs, as it suggests that multiple models could potentially describe the same underlying process. Especially for systems with limited prior knowledge, further investigation is needed to determine whether this flexibility captures biological variability or whether it points to the need for additional constraints to narrow down the solution space to the most biologically relevant model.

Despite its strengths, CLERA has some limitations that should be addressed in future work. First, the model’s performance is sensitive to hyperparameter choices, particularly the dimension of the latent space and library of candidate functions. Developing more robust methods for automatic hyperparameter tuning could improve the model’s consistency. Second, while CLERA can handle noise in the data to some extent, high noise levels can still impact its performance. Further improvements in denoising and data preprocessing techniques could enhance the model’s robustness. Third, the interpretation of the learned latent space and governing equations requires careful consideration. While we have shown that the model captures biologically relevant information, additional work is needed to develop standardized methods for interpreting these results in the context of specific biological questions.

In conclusion, CLERA represents a significant step forward in our ability to uncover the governing principles of complex biological systems from high-dimensional scRNA-seq data in a parsimonious and generalizable manner. By bridging the gap between data-driven discovery and mechanistic understanding, this approach has the potential to deepen our understanding of cellular processes and the interactions between active gene programs.

## Methods

### Autoencoders

An autoencoder is a neural network architecture consisting of an encoder and a decoder (46). The encoder compresses high-dimensional input data into a lower-dimensional latent representation reducing dimensionality while preserving essential information. The decoder reconstructs the original data from this latent representation. Regularizations and architectural choices can be applied to constrain the latent embeddings, influencing their quality and utility. This flexibility allows autoencoders to be used for dimensionality reduction, feature learning, and data denoising.

Training involves minimizing the reconstruction error, typically measured by mean squared error, to align the input with its reconstruction. For an input X, the encoder E produces latent embeddings z = E(x), which the decoder D uses to reconstruct the input as 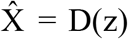. The reconstruction loss guides the optimization process so that the latent layer accurately captures the input data. The reconstruction loss for inputs is given by:

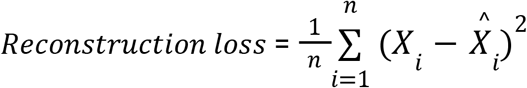

### SINDy and SINDy autoencoder

SINDy is a regression technique used to discover a best-fit dynamical system from training data (7). It takes input data *x*_*t*_ ∈ ℝ^*n*^ and predicts a system of ODEs as 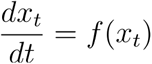, where *f* is a library of candidate functions composed of basis functions that we want SINDy to model. SINDy selects a subset of active terms from these candidates, promoting parsimony in the model, resulting in a simpler and more interpretable model that captures the essential dynamics. For *m* input snapshots, we stack them to form *X* = [*x*_1,_ *x*_2,_ …,*x*_*m*_]^*T*^ and compute the corresponding derivatives 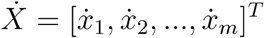. The library is constructed as Θ (*X*)=[1, *θ*_1_ (*X*), *θ*_2_ (*X*),…, *θ*_p_ (*X*)] ∈ ℝ ^*m×p*^, where *p* is the length of the candidate library, and *θ*_*i*_ are the candidate model terms. The coefficient matrix can be represented asΞ =[ *ξ*_1_, *ξ*_2_, … *ξ*_*n*_], where *ξ*_*i*_ contains the coefficients for the library functions of differential equation i.

To address the challenge of high-dimensional data, SINDy autoencoders leverage an autoencoder to find a low-dimensional state *z*= *ϕ*(*x*), where is *x* the time-series input, *z* is the latent embedding, and is the encoder (1). We incorporate sparse regression into the autoencoder training to facilitate the simultaneous discovery of latent embeddings and the corresponding ODE. The SINDy Autoencoder finds a latent dimension *z*_*t*_= *ϕ*(*x*_*t*_) ∈ ℝ^*d*^, where is *d* the size of the latent embedding, and an associated dynamic model 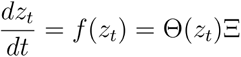, where *f* has few active terms.

We train the autoencoder with additional SINDy losses as regularizations to ensure that the latent embeddings have a parsimonious model representation. The reconstruction loss, 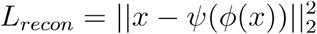 is used which ensures accurate embeddings, while the SINDy weight regularization, *L*_*SINDyReg*_= ||*ξ*||_1_ promotes sparsity of the coefficients. We also define two SINDy losses: 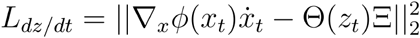, which enforces accurate prediction of dynamics, and 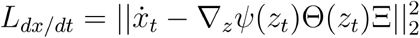, which reduces the error between the real-time derivatives of the input and the predicted derivatives using SINDy. We threshold the coefficients below a certain magnitude at regular intervals to promote parsimony. Finally, we conduct an additional refinement phase without *L*_1_ loss and thresholding for a few epochs to refine the coefficients.

### Overview of CLERA

CLERA is a deep learning-based model discovery method designed to uncover parsimonious and interpretable dynamical systems and active gene programs. It employs a specialized loss function to facilitate the joint discovery of a low-dimensional, interpretable latent space. The method’s architecture integrates multiple key components to analyze gene expression data. At its core, CLERA processes gene expression data to generate a cell-by-latent embeddings matrix. This latent representation serves as a crucial intermediate step, enabling three primary functions:

1. Data Reconstruction: A decoder network uses latent embeddings to reconstruct the original count matrix.
2. Cell-Type Classification: The latent space is used to classify distinct cell types within the dataset.
3. Dynamical System Discovery: CLERA incorporates SINDy to uncover a system of ODEs. These ODEs describe the temporal dynamics and interactions of the latent embeddings over the underlying biological processes.

This integration allows CLERA to simultaneously capture cell identities, gene expression patterns, and temporal dynamics.

Unsupervised learning of identifiable nonlinear representations has long been recognized as theoretically impossible without incorporating inductive biases or suitable constraints on the model (16). This fundamental challenge in representation learning has led researchers to explore various approaches to learn meaningful and causal representations of complex data. Autoencoders have emerged as a powerful tool for unsupervised representation learning, capable of discovering useful coordinate transformations and dimensionality reductions. However, while autoencoders can be trained in isolation, there is no guarantee that the learnt coordinates will be associated with sparse and physical dynamical models. To address this, CLERA integrates multiple components to learn interpretable and dynamically meaningful representations. By simultaneously training an autoencoder and a SINDy model, we constrain the network to learn coordinates associated with parsimonious dynamics. Also, CLERA incorporates a classifier connected to the latent embedding, leading to joint optimization of reconstruction, dynamics modelling, and classification tasks. By adding the classification task, the model can leverage additional information during training, to improve its ability to effectively capture the relevant dynamics of cell-specific behaviours. This integration introduces the inductive biases, potentially resulting in more meaningful and interpretable embeddings of the system, while outputs a system of ODEs that describe the interactions between these latent variables.

### Data Preprocessing

To preprocess the cell-by-gene matrix, the top 2,000 highly expressed genes are selected to focus on the most relevant features. The data is then normalized and log-transformed to stabilize variance and make the distributions of gene expression levels more comparable. The resulting matrix is analyzed using pseudotime (38) to establish a temporal sequence in the data and enable the conversion of static gene expression profiles into a dynamic time series which is essential for SINDy modelling. Given the sensitivity of SINDy to noise and the inherently noisy nature of scRNA-seq data, the time series is constructed by averaging gene expression counts over a sliding window of four-time steps (pseudo cell) with a stride of two steps. We experimented with multiple sliding window lengths but found that smaller window sizes would result in less effective denoising, while larger window sizes would lead to insufficient training data. Ultimately, a window length of 4 was chosen as the optimal balance between effective denoising and maintaining sufficient training data (Supplementary Note 1). This averaging process helps to smooth the data, reduce noise, and increase power. During this process, the cell type within each window is determined and assigned to ensure that cell identity is preserved. Derivatives of gene expression are calculated under the assumption of unit time steps between cells, a reasonable assumption given the continuous nature of pseudotime. This allows for consistent derivative estimation across the trajectory. Constructing the time series requires careful selection of cell types based on biological relevance, as some cell types may not contribute meaningfully to the dynamics being studied. This selection ensures that the resulting time-by-gene matrix accurately reflects the biological processes of interest

### Components of CLERA Objective function

The preprocessed time series results in a pseudo-cell by gene matrix. CLERA takes each step as input to the autoencoder and generates reconstructions of the time-averaged gene expression for that box. These steps form a batch, consisting of 1292 samples in the Pancreas dataset and 1172 samples in the hematopoietic differentiation dataset. During training, the derivatives calculated during preprocessing are used to calculate the SINDy losses while the cell type serves as the label for the classification network.

During classification, the latent representation is passed through a classification network, which outputs the probability of the embedding belonging to each cell type. The cell types are represented as labels *l* = [*l*_1_ *l*_2_ … *l*_*c*_ ] where c is the total number of cell types. The classification network *C* produces the probability of the input belonging to each cell type, *y* = *C*(*z*) where *y* = [ŷ_1_ ŷ_2_… ŷ_*c*_] and ŷ_*j*_ is the probability of the label being j. Multi-class cross-entropy loss 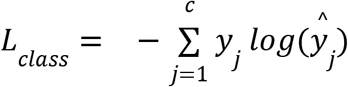 is used for this task. To improve stability during training, L1 regularization *L*_*NetReg*_ = ||φ||_1_ + ||ψ||_1_ + ||*C*||_1_ is applied to the network weights. This regularization encourages sparsity in the weights, reducing the risk of overfitting and improving the model’s generalization capabilities.

The loss function for CLERA is a weighted sum of 6 losses: Reconstruction Loss, Classification Loss, SINDy loss over the latent embedding, SINDy loss over the reconstructed genes, SINDy weight regularization and Network weight regularization which can be represented as

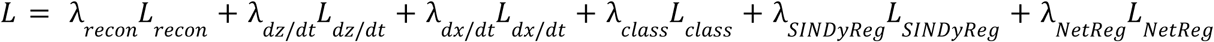

where different *λ*_*loss type*_ represent the loss weights.

### Modelling choices and training

The model hyperparameters significantly influence the nature of the resulting equations and require careful tuning to ensure effective training and accuracy of CLERA. Several conditions are essential for this process. First, we monitor the loss curves and the number of active SINDy coefficients closely, as they provide key insights that guide refinement. Second, the functional forms used in the model are designed to align with established prior knowledge of the process. Additionally, we make assumptions to maintain the integrity of the system’s dynamics: the right-hand side of every equation in the ODE should be non-zero, as a zero value would indicate that SINDy was unable to model the temporal features for that latent variable. Moreover, the discovered equations should be parsimonious, balancing expressiveness and simplicity. The dimension of the autoencoder’s latent layer plays a crucial role in this balance, which shows the trade-off between model complexity and parsimony. As the size of the latent dimensions increases, the SINDy library expands quadratically. Therefore, selecting the appropriate latent dimension is crucial, as a larger latent space can lead to more complex candidate functions, which complicates the model and increases the risk of overfitting. Experiments with higher-dimensional latent spaces (e.g., dimensions of 10 or 15) reveal that the model consistently produces ODEs where the right-hand side is zero for some of the latent variables. This indicates that higher latent dimensions produce variables that are less critical for capturing the dynamics of the system. Based on these findings, the final model uses a latent space of 6 dimensions, which provides a balance between capturing essential dynamics and avoiding unnecessary complexity (Supplementary Note 1). The autoencoder and classification weights are initialized with Xavier initialization which is known to promote stability and efficient learning. We train both the pancreas and bone marrow datasets each for 50 times and we select the best model and parameters considering the conditions and criteria above such as the lowest loss, parsimony, and model complexity.

In modelling complex biological systems, it is crucial to leverage the prior knowledge we have of the system to carefully select a library of candidate functions. This selection should include functions capable of capturing a wide range of dynamics. Studies have considered various terms, such as polynomials up to degree 3, Michaelis-Menten kinetics, fractional terms, and exponential functions, to model nonlinear and dynamic processes (47–51). Building on these approaches, we carefully select our SINDy library to include linear, second-order polynomial terms along with exponential, sinusoidal and fractional function forms. The sinusoidal terms are crucial for capturing oscillatory behaviours, while the exponential terms effectively model processes involving growth or decay. The fractional function helps with modelling nonlinear behaviours typically observed with biological processes, such as Michaelis-Menten kinetics. The SINDy model coefficients are randomly initialized with 1 and -1 which ensures that there is no bias towards one particular direction (positive or negative coefficients). This makes the model better at discovering the true underlying relationships in the data. Refer to Supplementary Table 1 for different Hyperparameters that are chosen while training CLERA on pancreas and bone marrow datasets.

For the bone marrow dataset, the autoencoder weight and SINDy coefficient initialization are done in a transfer learning framework. The SINDy library and autoencoder architecture are consistent across both the pancreas and bone marrow, ensuring that the trained weights and SINDy coefficients of the pancreas can be used as initialization for bone marrow. CLERA is initially trained on the Pancreas dataset using 15 different random initializations, resulting in 15 distinct instances of trained Pancreas weights. These trained instances are then evaluated on the bone marrow data. The top 6 experiments that exhibit the lowest losses when tested on the bone marrow data are selected. From these 6 experiments, an average of the Autoencoder weights and SINDy coefficients is computed. This averaged set of weights and coefficients is used as the initialization for subsequent bone marrow training. By leveraging transfer learning in this manner, the model benefits from faster convergence and improved performance, as the pre-trained weights provide a more informed starting point (Supplementary Fig 4).

Another key difference between the pancreas and bone marrow training lies in the SINDy thresholding interval. For the bone marrow dataset, the thresholding interval at 500 epochs is set higher than that for the pancreas dataset which is kept at 150. This adjustment is necessary as the SINDy coefficients derived from the pancreas experiments exhibit magnitudes much lower than 1, due to the application of L1 regularization. By setting a higher threshold for the bone marrow training, the network is better at adjusting and correcting the coefficient magnitudes, which ensures that key SINDy terms do not undergo early thresholding.

In the unsupervised training setting, where there is an absence of explicit target ODE terms and structure, an inductive bias is introduced to determine the appropriate number of training steps. A maximum acceptable term threshold is established, defining the maximum number of active SINDy coefficients permitted on the right-hand side of the ODE. During training, once the number of active coefficients falls below this threshold, the refinement phase is initiated, consisting of an additional 150 epochs. This threshold is kept at 30 to ensure that the model remains parsimonious.

### Personalized Pagerank

PageRank is an algorithm that ranks nodes in a graph by evaluating the number and weight of links between them. Nodes that are connected to highly-ranked nodes are deemed more important and thus receive a higher PageRank score. The standard PageRank algorithm calculates a global importance score, treating all nodes equally in the process.

Personalized PageRank, on the other hand, is an extension of this algorithm that computes node rankings relative to a specific node or set of nodes within the graph. Unlike traditional PageRank, which provides a uniform importance score across the entire graph, PPR introduces a bias toward the selected nodes. This bias allows for the calculation of personalized importance scores, emphasizing the relevance of nodes in relation to the chosen node(s) of interest.

The interaction graph is constructed by including the gene-latent variable interaction using the top “K” SHAP for each latent variables and inter-latent-variable interaction via the ODE. As cells progress further in pseudotime and become more dissimilar, the mean intersection percentage between top-k genes of different latent variables decreases for all latent variables (Supplementary Fig 6, Supplementary Note 2). To rewire the original interaction graph and capture direct gene interactions, we use PPR to quantify the probability of connectivity between genes. This approach models the likelihood that a random walk starting from one gene will visit another, thereby capturing both direct and indirect relationships in the network.

For each gene i, we initialize a personalized vector *v*_*i*_ where the ith entry is set to 1 and all other entries are 0. This biases the random walk to originate from gene i. We then compute the PPR vector π_*i*_ by recursively solving the equation:

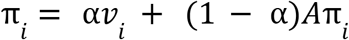

where A is the column-normalized adjacency matrix of the initial network and α is the teleportation probability (typically set to 0.85).

The resulting α_*i*_ vector contains PageRank scores for all genes relative to gene i. We interpret these scores as probabilities of connectivity, with higher values indicating stronger relationships. To construct the refined network, we select the top “E” edges for each gene based on these probabilities. In our implementation, we set E=600 to balance network sparsity and connectivity (Supplementary Note 2). This process is repeated for all genes, resulting in a directed, weighted graph that emphasizes the most significant regulatory relationships in the network. Also, the PPR-based rewiring captures higher-order network topology beyond direct interactions, which can reveal important indirect regulatory connections.

Calculating various centrality metrics on the obtained connectivity graph helps us identify the most influential genes, determine hubs of activity, and uncover critical pathways. Each centrality metric offers a unique perspective on the role and significance of nodes in the network. The centrality metrics used include:

- **Degree Centrality**: Degree centrality measures the number of direct connections a node has in a network. Nodes with a high degree of centrality are typically more active or prominent within the network.
- **Closeness Centrality**: Closeness centrality assesses the average length of the shortest paths from a node to all other nodes in the network. Lower scores indicate that the node occupies a more central and important position in the network.
- **Eigenvector Centrality**: Eigenvector centrality evaluates a node’s influence based on both the quality and quantity of its connections. A node is more central if it is connected to other central nodes, highlighting its importance within the network.
- **PageRank**: PageRank assigns importance to a node based on the principle that connections from high-ranking nodes contribute more to a node’s score, making it a more influential node in the network

By analyzing these centrality metrics, we identify the key genes in the interaction network, gene markers and hubs that can play a key role in cell function and gene interaction.

### Simulated data

To evaluate SINDy’s ability to detect governing equations, we design a simple two-gene regulatory network. This system is governed by stochastic differential equations, incorporating both deterministic dynamics and stochastic fluctuations. The network features two mutually inhibiting genes, described by *G*_1_ and *G*_2_:

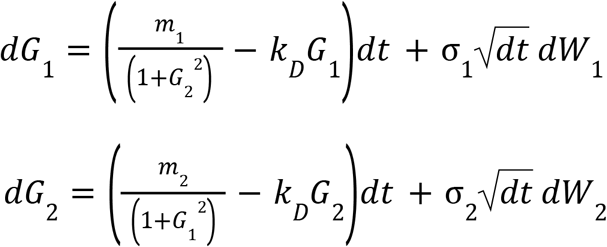

In this model *k*_*D*_ represents the degradation rate, *m*_1_ and *m*_2_ determine the synthesis rates of the genes, σ_1_ and σ_2_ are noise intensities, *d*t is the time step, *dw*_1_ and *dw*_2_ are increments of independent Wiener processes modelling Gaussian noise. By adjusting these parameters, the system can exhibit saddle-node and pitchfork bifurcations. To generate training data for SINDy, we set *m*_1_ =3, *m*_2_ =1, *k*_*D*_ =1 and σ_1_ = σ_2_ = 0.2. The Euler–Maruyama method was used to simulate these stochastic differential equations, to capture the system’s dynamics while accounting for noise that scales with time steps.

## Data availability

The datasets used in this study are publicly available in public repositories. Simulated scRNA data was generated using SERGIO, which is accessible at https://github.com/PayamDiba/SERGIO. Preprocessed versions of both the pancreas development data (accession number GSE132188) and the human hematopoiesis data (accessed through the Human Cell Atlas data portal under the Human Hematopoietic Profiling project) can be downloaded from https://scvelo.readthedocs.io/en/stable/.

## Code availability

Our Python implementation of CLERA can be found at:

